# Comparison of manual and machine learning image processing approaches to determine fungiform papillae on the tongue

**DOI:** 10.1101/2020.07.05.187658

**Authors:** Camilla Cattaneo, Jing Liu, Chenhao Wang, Ella Pagliarini, Jon Sporring, Wender L.P. Bredie

## Abstract

Human taste perception is associated with the papillae on the tongue as they contain a large proportion of chemoreceptors for basic tastes and other chemosensation. Especially the density of fungiform papillae (FP) is considered as an index for responsiveness to oral chemosensory stimuli. The standard procedure for FP counting involves visual identification and manual counting of specific parts of the tongue by trained operators. This is a tedious task and automated image analysis methods are desirable. In this paper a machine learning image processing method based on a convolutional neural network is presented. This automated method was compared with three standard manual FP counting procedures using tongue pictures from 132 subjects. Automated FP counts, within the selected areas and the whole tongue, significantly correlated with the manual counting methods (all ρs ≥ 0.76). When comparing the images for gender and PROP status, the density of FP predicted from automated analysis was in good agreement with data from the manual counting methods, especially in the case of gender. Moreover, the present results reinforce the idea that caution should be applied in considering the relationship between FP density and PROP responsiveness since this relationship can be an oversimplification of the complexity of phenomena arising at the central and peripherical levels. Indeed, no significant correlations were found between FP and PROP bitterness ratings using the automated method for selected areas or the whole tongue. Besides providing estimates of the number of FP, the machine learning approach used a tongue coordinate system that normalizes the size and shape of an individual tongue and generated a heat map of the FP position and normalized area they cover. The present study demonstrated that the machine learning approach could provide similar estimates of FP on the tongue as compared to manual counting methods and provide estimates of more difficult-to-measure parameters, such as the papillae’s areas and shape.

## Introduction

The ability to detect and differentiate between food-derived chemical stimuli is mediated by receptor cells within taste buds [1], which primarily reside within the tongue taste papillae (fungiform, foliate and circumvallate). Among them, the fungiform papillae (FP) are the anatomical structures foremost involved in the detection and transduction of oral and somatosensory stimuli. Given the double innervation of FP, these anatomical structures has been selected as one of the phenotypic markers of taste sensitivity, due to their relative abundance and accessibility on the tongue anterior part, their association with the density of taste buds [2,3] and the higher inter-individual variability in their number and shape among human subjects.

Moreover, Several studies have reported on the positive correlation between the density of FP and the perceived intensity of a the bitter substance 6-n-propylthiuracil (PROP) [4–8], the other extensively accepted phenotypical marker of taste responsiveness. Indeed, subjects characterized by an higher responsiveness to this compound, i.e. the *PROP super-tasters*, present at the same time a high number of FP [5]. However, more recent studies, involving large population samples, have failed to find a relationship between the density of FP and responsiveness to chemosensory stimuli [9,10]. These contradicting results could be due to issues related to the FP-identification and -counting methodology [11–13].

Video-microscopy was the first non-invasive technique used to assess the density of FP in a clinical setting [2,3,14]. However, this method has several disadvantages, since it is difficult to apply outside of a clinical setting and takes up to 60 minutes to collect images [14–16]. A suitable substitute for video-microscopy is the digital photography, the most recent and widely used technique for FP density assessment [17]. This method is based on the visual inspection of high-resolution digital pictures of blue-stained tongues [16], followed by manual counting by operators trained according to the Denver Papillae Protocol guidelines [11]. The adoption of this standardized protocol can improve consistency among operators, but not fully remove the bias associated with the papillae counts [18]. Thus, visual inspection of digital pictures remains subjected to researcher bias and it is restricted to specific areas of the tongue. Furthermore, manual counting is limited in other parameters of the papillae such as the papilla shape, surface area and density across other parts of the tongue.

Recently, automated methods to detect FP from digital images have been proposed [12,17,19,20], demonstrating the increasing interest in this issue. However, most of these methods still have some limitations related to i) their raw counting accuracies; ii) requiring images to be taken under very specific conditions; and iii) limited robustness to papillae with non-typical appearance.

A more versatile method for analysis of digital tongue pictures would be the alignment in position and projection onto a standardized fixed tongue in analogue to brain imaging analysis. On this basis, we suggested the use of convolutional neural networks based on state-of-the-art deep learning. These approaches do not depend on the author’s manually engineered features, but instead mathematically learn how to extract, understand, and generalize the most distinguishing features that define a papilla. Based on this assumption their segmentation results could be much more accurate and more robust to varied and even previously unseen data.

Specifically, the present study describes a novel automated procedure for counting and evaluating FP based on state-of-the-art deep learning. The relationships between automated method response and manual counting were investigated. FP distribution greatly varies depending on the area of the count (e.g. FP on the tongue tip will tend to be higher in number than those counted in more posterior tongue regions). Thus the present paper compared, for both manual and automated procedures, the counts derived from the area located at 1 cm from the tongue tip along the median line, considering the differences in terms of area and shape (circular or squared). Moreover, the final goal is to improve papillae segmentation to such an extent that we will not only be able to get more reliable counts, but also open up the possibility of using other difficult-to-measure parameters, such as the papillae’s areas and shape, for future research.

## Results

### Descriptive statistics of manual and automated counts

The distribution of FP across the 132 subjects using the manual counts tended all to a non-normal distribution (Manual count in ED1: W = 0.97, p < 0.05; Manual count in ED2: W = 0.95, p < 0.001; Manual count in ED3: W = 0.96, p < 0.001), with data skewed to the right. Before any analysis related to automated counting was performed, we went through the images and removed 20 images with major quality issues. The distribution of FPD across the 112 subjects using the automated counts in the selected area tended all to a normal distribution (Automated count in ED1: W = 0.99, p = 0.62; Automated count in ED2: W = 0.99, p = 0.23, Automated count in ED3: W = 0.99, p = 0.32).

Descriptive statistics of manual and automated FP counts are reported in **Table 1.**

**Table 1.** Descriptive statistics of papillae density from manual count (Manual ED1-ED3) and automated image analysis (Automated ED1-ED3).

| Descriptive statistics | Manual |  |  | Automated |  |  |
| --- | --- | --- | --- | --- | --- | --- |
|  | ED1<sup>a</sup> | ED2<sup>b</sup> | ED3<sup>c</sup> | ED1 | ED2 | ED3 |
| Min | 8 | 6 | 31 | 7.3 | 5 | 27 |
| Max | 32.3 | 32.5 | 142 | 32.0 | 30.5 | 123 |
| Mean | 17.6 | 14.5 | 72.6 | 17.3 | 15.4 | 75.5 |
| Standard deviation (Std) | 4.9 | 4.9 | 20.0 | 4.2 | 4.4 | 17.2 |
| PROP status |  |  |  |  |  |  |
| NT<sup>d</sup> (mean ± Std) | 16.3 ± 4.3 | 13.5 ± 4.6 | 67.9 ± 18.7 | 17.8 ± 3.2 | 16.0 ± 3.7 | 78.5 ± 15.2 |
| MT<sup>e</sup> (mean ± Std) | 17.1 ± 4.9 | 14.1 ± 5.0 | 70.7 ± 18.9 | 17.1 ± 4.3 | 15.3 ± 4.3 | 75.1 ± 16.9 |
| ST<sup>f</sup> (mean ± Std) | 19.0 ± 5.2 | 15.5 ± 4.9 | 77.7 ± 21.5 | 17.3 ± 4.7 | 15.2 ± 5.0 | 74.6 ± 18.8 |

<sup>a</sup> Counting method according to Nachtsheim and Schlich [7]
<sup>b</sup> Counting method according to Masi et al. [30]
<sup>c</sup> Counting method according to Essick et al. [5]
<sup>d</sup> Non-tasters; <sup>e</sup> Medium tasters; <sup>f</sup> Supertasters

### Comparison between manual and automated counts

A comparison between automated and manual counts within the selected areas and the whole tongue (TOT) has been performed through Spearman’s correlation analysis (see **Table 2**).

**Table 2.**
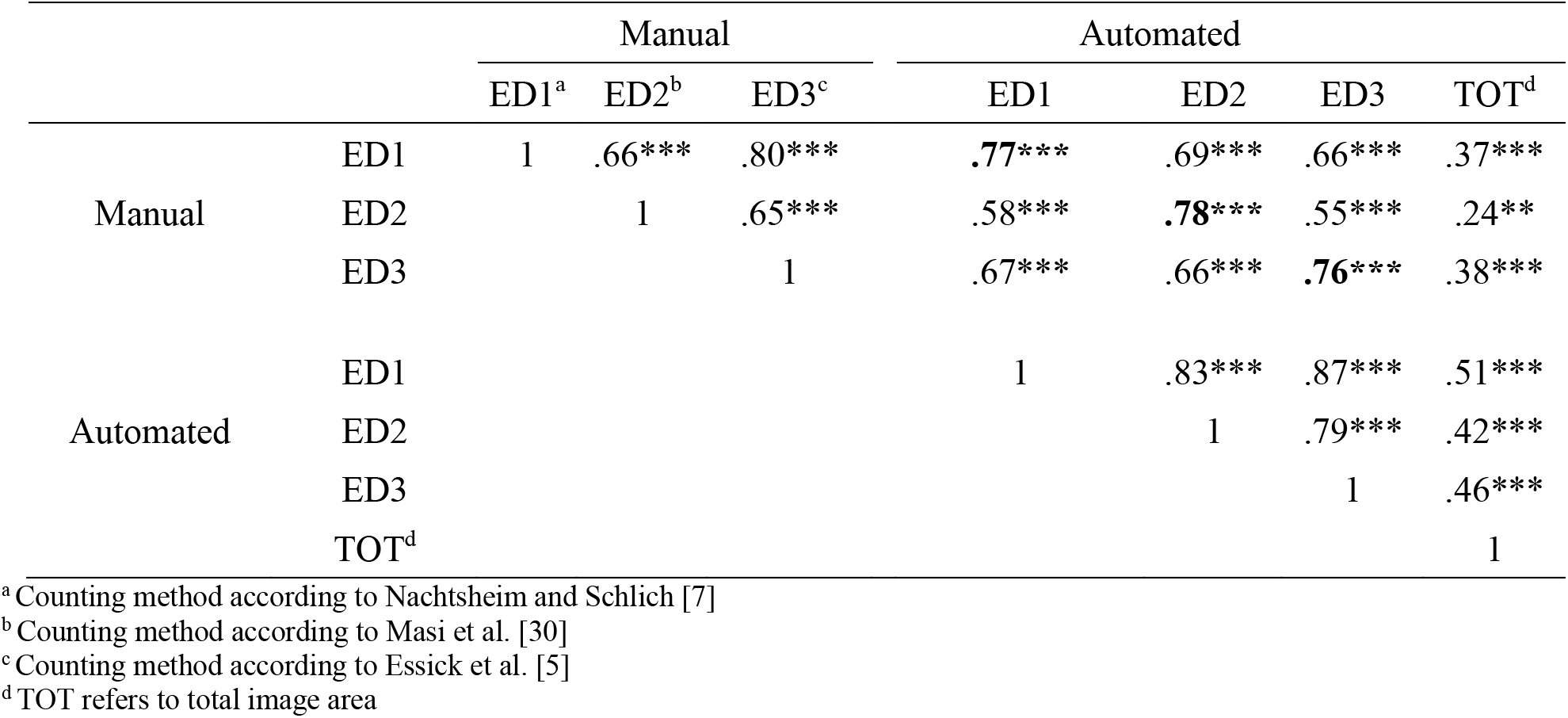
Correlations among manual and automated counts: spearman’s correlation matrix (n =112). Values in bold represent significant correlations between manual and its respective automated count in each selected area.

The correlation between the manual and automated counts has also been evaluated by plotting them against each other (see **Fig. 1**).

**Fig. 1.**
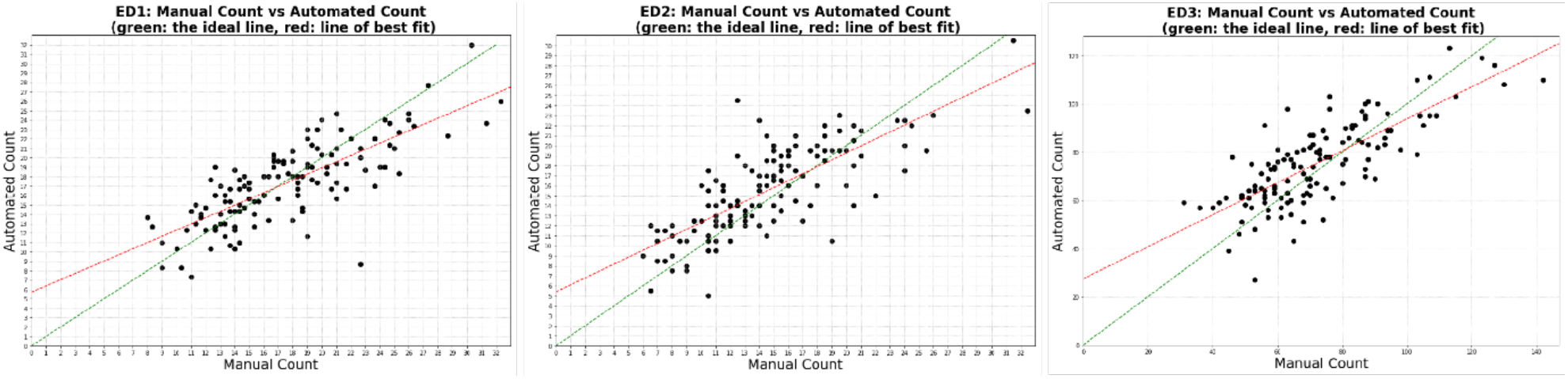
Automated counts plotted against manual counts in selected areas (ED1–3).

The green and red lines in each subplot of **Fig. 1** represent the ideal line (*y* = *x*) and the linear regression respectively. Together, they showed a clear visual correlation between the automated and manual counts for all three regions. A similar trend can be found when looking at the spearman’s coefficients in each region, with ED3 being the lowest at ρ_s_ = 0.76, ED1 in the middle at ρ_s_ = 0.77, and ED2 the highest at ρ_s_ = 0.78.

The correlations among taste function phenotypic measures (PROP and FP counts, both manual and automated) were also tested. PROP bitterness ratings were positively correlated to the manual counts in ED1-ED3, albeit the magnitude of this association was very low (in all selected area ρ_s_ <0.2, p<0.05) whereas no significant correlations were found between FP automated counts, neither the whole tongue nor PROP bitterness ratings.

An objective-based evaluation of our model has been made, where we compared its ability to distinguish between different PROP taster status classes with that of the manual counts (non-tasters, NT; medium tasters, MT, supertasters, ST). To check if the manual counts and automated counts are detecting the same pattern, we first plotted the distributions of mean FP counts computed from bootstrapping each PROP taster status class in **Fig. 2.**

**Fig. 2.**
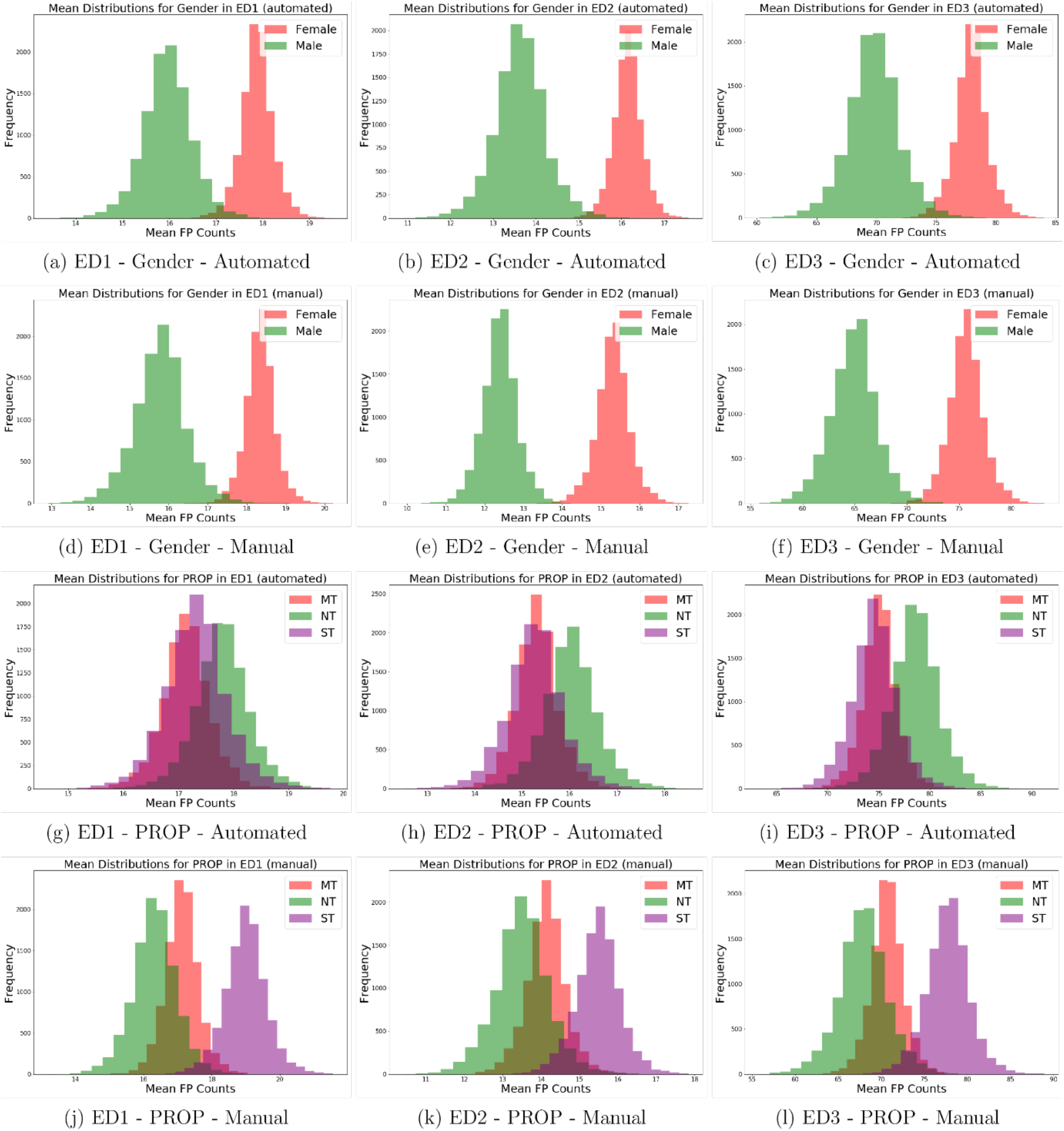
The bootstrapped mean distributions of PROP taster status classes.

The results showed that manual and automated counts detected the same patterns for gender but are not in full agreement for PROP taster status.

In the case of gender, a higher mean FP count was consistently observed in females than males by both counting methods across all selected areas. Judging from the medians of the mean distributions for gender, the automated model seemed to systematically overcount by approximately 1 – 2 papillae in ED1 and ED2 and by 4 papillae in ED3 as compared with the manual counts.

Regarding the relationship between PROP taster status and FP, the manual method detected higher mean FP counts in supertasters than non-tasters, whereas the automated method failed to detect any difference between the two groups, as shown by the significant overlap between their mean distributions. Upon closer inspection of the median values of the mean distributions, we can see that in ED2 and ED3, the automated counts for supertasters match the manual counts quite well, whereas medium tasters and non-tasters were slightly overcounted by the automated method, especially for ED1. The weakly contradicting conclusions reached for PROP non-tasters and supertasters are thus caused by our model’s tendency to overcount FP in non-tasters as compared to manual counting. Several counting experiments were carried out to obtain a more detailed understanding of why this happened. We started by recounting a randomly sampled subset of non-tasters and supertasters to check the precision in manual counting (**Fig. 3 a-b**).

**Fig 3 a-b.**
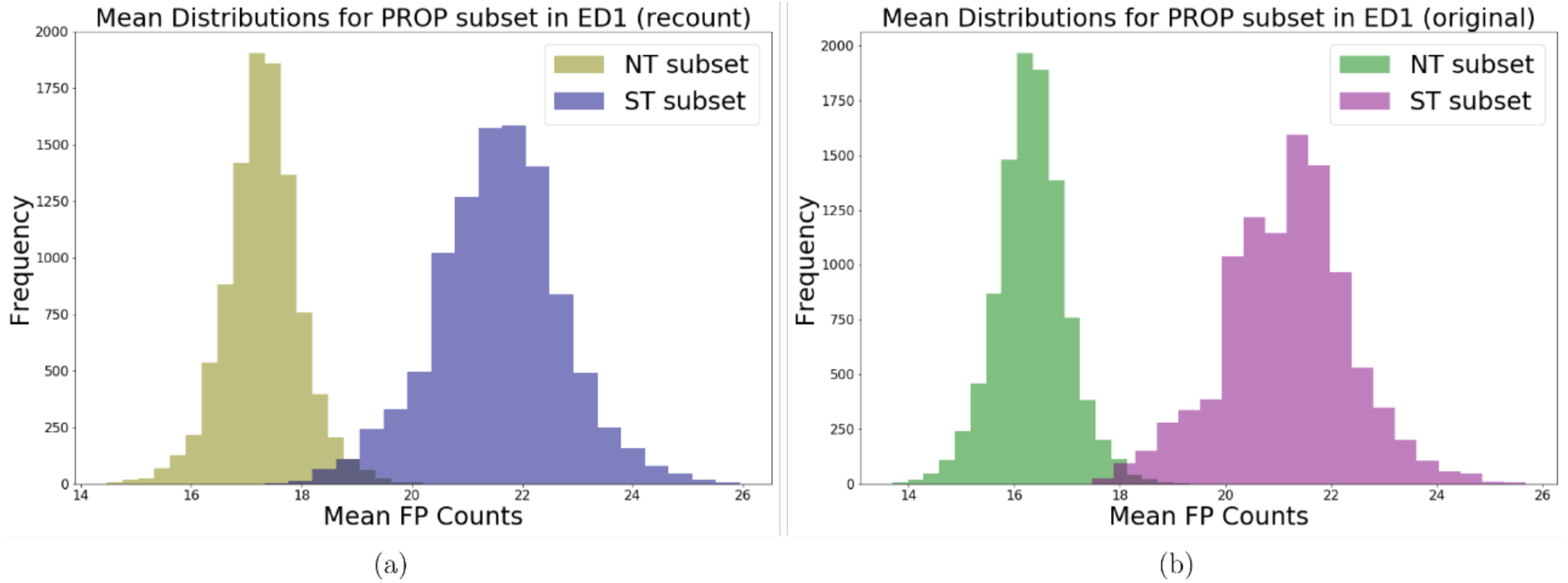
Bootstrapped mean distributions in ED1: (a) is computed using a recounted subset of PROP and (b) is computed from the same subset’s original manual counts.

The recounting of subsets of non-tasters and supertasters has shown that while the precision is quite high for supertasters, the mean distribution for non-tasters shifted right by 1 papilla in the recounting. Although this does not change the conclusion reached by the manual method, it does suggest that the actual difference between non-tasters and supertasters may not be as large as previously shown in Figure 6.

To test the statistical significance of the observed difference in mean FP count between females and males, a bootstrapped permutation test was performed, and its results are plotted below. Given uncertainties associated with the PROP results, we would only consider the gender category from this point on.

As shown in **Fig. 8 (a-c)**, both automated and manual counts were able to reject the null hypothesis in all three regions of interest. However, the permutation test results of manual counts are still more accurate (i.e. presents lower variability) than that of the automated counts.

**Fig. 4 a-c.**
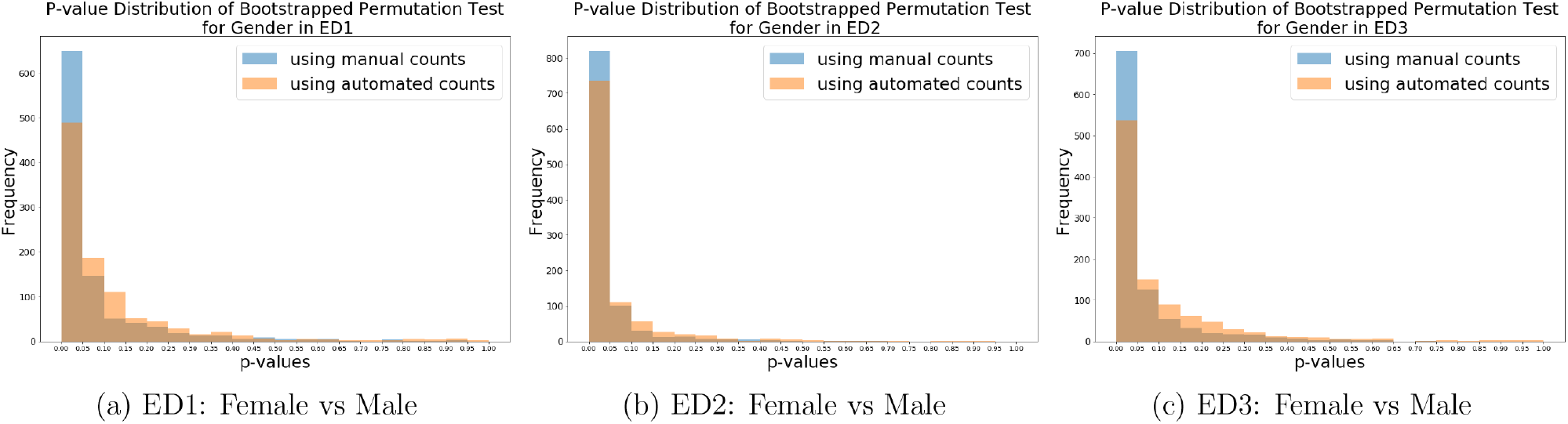
p-value distributions of bootstrapped permutation test on the mean FP counts for gender.

## Discussion

The present study describes a novel automated procedure for counting and evaluating the number of fungiform papillae (FP) based on state-of-the-art deep learning. The determination of the number of FP is an important measure in sensory science research since this measure has been used as index of taste sensitivity in general.

The majority of studies in the literature manually count the number of FP in a small region on the anterior tongue tip. Recently, automated methods to detect FP from digital images have been proposed. Sanyal and colleagues [12] proposed the first attempts at using computers to automatically analyze tongue papillae. The authors designed an algorithm using the TongueSim suite of software, which allowed them to measure the FP density and other properties, such as the degree of roundness of each papilla. The algorithms proposed in that study appeared generally robust in the identification and counting of FP. However their algorithm seemed tp present poor prediction in the lowest range of papillae counts, probably due to the very few number of pictures (n=9) used to validated the model. Valencia and colleagues [20], designed an algorithm that allowed the user to manually select a rectangle that contains an average papilla and then uses it as a template to find similar objects in the search area. The algorithm uses the 2D normalized cross-correlation between the greyscale versions of the tongue image and the template. When comparing the algorithm’s counts with manual counts, they were able to show a general correspondence between the two, where tongues with higher automatic counts also have higher manual counts. However, the proposed algorithm was very dependent on the user’s ability to provide a good exemplar FP to be used as a reference for the total FP count on the whole tongue. More recently, two different automated approaches were proposed for fully automated detection and calculation of the number of FP from digital images [13, 19].

Piochi and colleagues [13] proposed an algorithm based on the procedure used by Kraggerud et al. [21]. A script was developed to provide a black and white image through correction of the background variation and graphical emphasis of the elevated structures represented as white spots. Then, the procedure allowed to automatically identify the FP as circular-like elements (whit a diameter ranging from 0.30 to 1.05 mm) and to compute the frequency of these circular-like elements in classes with varied diameter size. Eldeghaidy and colleagues proposed a color-based segmentation method based on the procedure used by Rios et al. [22]. The authors manually selected three structures of interest – fungiform papillae, filiform papillae and tongue base, and transformed the colour space of these regions from RGB into LAB to minimize the sensitivity for illumination differences, before taking the average colour of each region. The segmentation works by taking the nearest neighbour according to the Euclidean colour separation between the regions of interest and each pixel. However, these last two different models cannot be compared directly unless the tongue images are of the same kind, e.g., all stained or all unstained, and have an equal quality standard. Nevertheless, for the case-specific performance, the count accuracies seem to have improved significantly since the previous attempts [12, 20].

The novel automated procedure presented, applying the deep learning approach, had the purpose of achieving more accurate and comprehensive outcomes compared to the existing works within this area. Firstly, deep learning has been incorporated into the segmentation stage, allowing the model to automatically learn the features that best describe a papilla and how to interpret them, thereby resulting in more accurate segmentation. Moreover, the high-performance of the computerized approach permitted the computation of the pixelwise distribution with normalized convolution, in order to minimize information loss as much as possible.

In general, results from the automated model matched those from the manual count in each selected area. The results showed a strong agreement with manual counting in all the selected areas (all ρ_s_ ≥ 0.76), as well as the overall FP count on the anterior part of the tongue strongly correlates with the automated counts.

An in-depth comparison between what the model and the expert’s counts has also been made. However, the classification of fungiform papillae was sometimes difficult for both model and expert due to the FP 3D shapes (they cannot be flat) and image issues associated with blurriness and poor staining. The model was systematically slightly more lenient on papillae inclusion in comparison with the expert, causing a systematic overcounting.

Indeed, according to Nuessle and colleagues [11], the Denver Papillae Protocol criteria to identify and, thus manually count, a tongue structure as FP is to strictly assume a circular shape and a diameter size of at least 0.5 mm. Thus, computer models could identify as FP tongue structures with irregular shapes or with a wider variety of sizes. This could result in an underestimation or, contrarily, in an overestimation of structures with a certain size/shape configuration, treating these elements as actual papillae and not as artefacts.

Moreover, for smaller analysis regions ED1 and ED2, the boundary effect is also in play – it was decided that an FP will be included in the counts if its centre point lies within the region, but judging this when counting manually proved to be difficult for FPs that are centred near the boundary. Despite of the randomness of this effect, we cannot exclude that this might have coincidentally resulted in the lower manual FP counts for non-tasters in Figure 6. This hypothesis has been corroborated by the recounted subset in Figure 7 suggesting that a representative selection of the initial dataset of pictures could improve the accuracy of the automated counting.

Apart from these discrepancies, our model showed in general good reliability (similar mean values among ED1-ED3 in automated and manual counts) comparing the two methods of counting. Nevertheless, the recounting of subsets of non-tasters and supertasters seems to have improved also the model accuracy (i.e. same elements counted by the different methods).

Irrespective of the imbalance in male/female ratio (males <25%) in our study, the females had a persistent higher FP count by both counting methods. Gender differences on the variations in taste functions have been recently reviewed [17, 23], suggesting that women detect basic taste stimuli at lower concentrations than men. However, whether these differences between sexes are reflected in the number of FP is still unclear and several studies failed to find a significant effect of gender [8,15,24–30]. On the contrary, other recent large-scale studies reported that women consistently having a higher number of FP in respect to men [9,10], and Piochi and colleagues [31] hypothesized that males and females may differ in the number of FP having specific diameters.

The relationship between FP count and PROP responsiveness was also assessed on both manual and automated counts, considering selected areas and the whole tongue. Significant correlations between manual counts of FP and PROP intensity ratings were found, albeit the magnitude of this association was very low (ρ_s_ <0.2), accordingly to some previous findings [4,7,8,32]. However, neither the number of FP measured with the automated counts in the selected area nor the total number of FP on the whole tongue significantly correlated with the PROP intensity ratings. Recent studies involving a large sample size of individuals failed to show an association between FP count and PROP rating (see [17] for a review). Moreover, it has been reported that subjects with lower FP density are characterized by increased sensory responsiveness [10,33]. Indeed, the present automated data showed that FP variation is slightly associated with the subject’s PROP taster status, due to significant overlap between the mean FP distributions. These distribution patterns seem to be in line with one found by Dinnella and colleagues [10], who explored the importance of FPD in taste sensing in a group of PROP NT and PROP ST. The authors showed that both ST and NT groups are characterized by individuals with low and high FP density, and hypothesized that the variation of density of FP strongly affected the orosensory perception of food stimuli with the same PROP taster group (i.e. high/low FP density in NT or ST subjects). Thus, as already concluded by the authors, additional insight should be gained on associations between FP/PROP, and the role of peripheral sensing organs should be reconsidered.

Although the deep learning model used in this paper succeeded in detecting most of the papillae, some consideration must be made in terms of its strengths and weakness. The biggest advantage of using a convolution neural network is that its classification rules are learned through examples rather than programmed. This generally leads to a more accurate segmentation, which can be further improved through the correction and relearning of mistakes. Boosting the performance of non-machine learning-based methods, on the other hand, will likely require significant design changes and will thus be much more difficult and time-consuming. A weakness of deep learning method is ironically also its reliance on learning from data. For the model to learn the right classification rules, a large number of highly accurate examples must be provided in the training set.

Considering the current state of automatic papillae analysis in this paper and related works, there is still further issues to be addressed. For instance, our counting experiment revealed that an object’s 3D shape also matters in the sense that active papillae cannot be flat. Given the difficulties faced by tongue experts and our model in judging this feature consistently, the current method of data collection should be improved. While problems associated with blurriness can easily be corrected by improved camera setup, other underlying issues do not have such a simple solution. This includes, but is not limited to, the dye adhesion problem in tongue staining and the very fact that we are trying to evaluate 3D shape using 2D images. It could be therefore worthwhile to add laser scanning to the existing data collection procedure or use a multi-camera system for stereoscopic reconstruction. Moreover, to eliminate possible boundary effects associated with manual counting in future studies, an FP should only be counted if it is entirely inside the analysis region.

Another area of the analysis worth exploring in future studies is how the segmentation masks can be used more effectively. At the moment, the papillae segmentations are converted into counts before any statistical tests are performed. As a result, many of the new segmentation features that can bring new perspectives into the study of fungiform papillae are overlooked (such as area and shape).

In conclusion, data from the present study demonstrated how a computerized approach, based on state-of-the-art deep learning, can open a whole new range of papillae features to looking into. The density of FP predicted from automated analysis output is in good agreement with data from the manual count, especially in the case of gender. Moreover, a significant improvement in the accuracy has been obtained through the recounting of subsets of non-tasters and supertasters. The deep learning machine approach for tongue analysis could provide further and innovative advantages by defining a tongue coordinate system that normalizes the size and shape of an individual tongue and by defining a heat map of the FP position and of the normalized area they cover. This could open up the possibility of using other difficult-to-measure parameters, such as the papillae’s areas and shape. The present study showed a prove of concept for automated papillae counting using a deep learning computer approach. The automated method appears to be suitable for FP counting in larger scale lingual surface imaging studies.

## Material and methods

### Subjects

One hundred and fifty-two subjects (23% males) between the ages of 18 and 55 years were recruited to attend the test and sensory and tongue picture data were collected anonymously. All subjects participated by informed consent. The study protocol was reviewed by the Danish National Committee on Biomedical Research Ethics (ref. nr: 18012023).

### Taste responsiveness to PROP

The protocol applied to evaluate subjects’ PROP responsiveness was fully described in Cattaneo et al. [34] using a method proposed by Prescott and colleagues [35]. Subjects were asked to rate the intensity of bitterness of a supra-threshold 0.0032 M solution of PROP (European Pharmacopoeia Reference Standard, Sigma-Aldrich) through the Generalized Labeled Magnitude Scale (0–100), gLMS [36]. Two identical samples (10 ml) were presented to the subjects who were asked to hold in their mouth each sample for 10 s, then to expectorate the solution and to evaluate the perceived bitterness intensity after 20 s. To avoid carry-over effect, subjects were asked to rinse the mouth with water after the first sample evaluation and to wait 90 seconds before evaluating the second sample [37]. The PROP sensitivity score for each individual was calculated as the mean of both gLMS measurements.

### Acquisition of tongue pictures

To measure the FP count, subjects were asked to place their chin on a chin rest table fixture (Western Opthalmics, USA) and extend their tongues and hold it in steady position. The tongue was stained with a blue food coloring using a cotton-tipped applicator. In this way, the FP became more clearly visible, as mushroom-like structures, on the anterior portion of the dorsal surface of the tongue. Digital pictures were recorded using a 18 megapixels digital camera (Canon EOS 700D, Japan) in a brightly lit room using the camera’s macro mode with no flash. Pictures captured the anterior part of the tongue. A ruler fixed on the chin rest provided a size calibration to allow surface comparisons across subjects. A series of pictures were taken and for each subject the best photograph was selected. Photoshop software (Adobe, USA) was used to mark the area to count the FP.

### Manual FP counting procedures

Three different counting methods were employed on the tongue pictures. These encompassed all FP counting at the tip of the tongue, but differed in the surface area and uni/bi laterality used for counting. The counting methods used were according to:

i. Nachtsheim and Schlich [7], in which three circles of 6 mm diameter were drawn in the front of the anterior (ED1; **Fig. 5a**);
ii. Masi et al. [30], in which two circles of 6 mm diameter were drawn one on the left side and one on the right side of the tongue, 0.5 cm from the tip and 0.5 cm from the tongue midline (ED2; **Fig. 5b**); and
iii. Essick et al. [5], in which a square of 1 cm^2^ was drawn on the tongue tip with the midline of the tongue bisecting the center of the square (ED3; **Fig. 5c**).

**Fig. 5 a-c.**
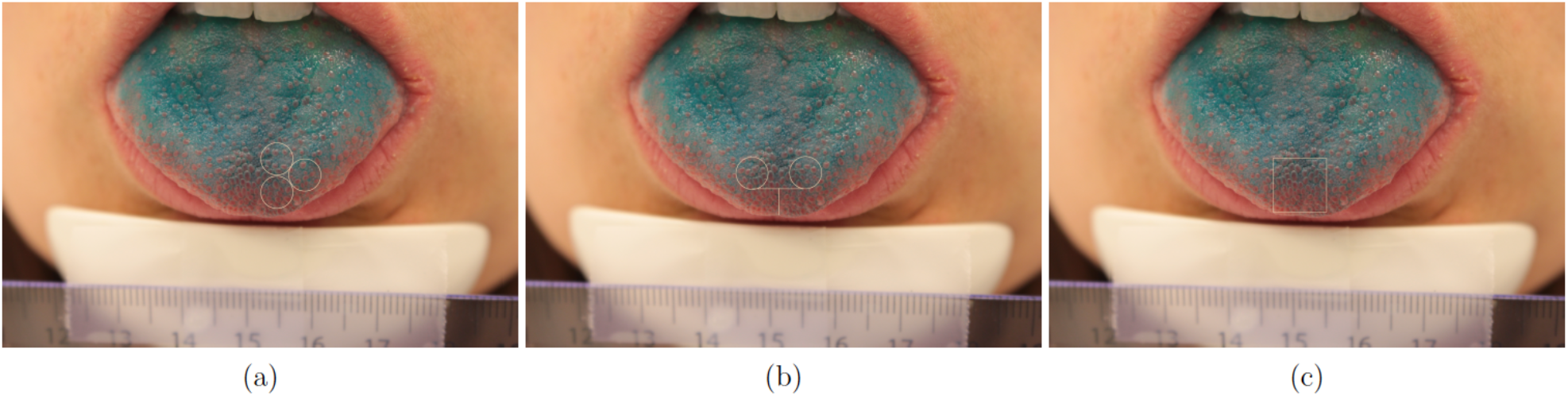
An example of a tongue image used for fungiform papillae counting (FP). The three counting areas shown in the subfigures are selected according to a) Nachtsheim and Schlich [7]–ED1; b) Masi et al. [30] – ED2; c) Essick et al. [5] – ED3.

For each method, the individual FP numbers were manually calculated following the Denver Papillae Protocol [11].

### Automated counting procedure

Before any analysis related to automated counting was performed, we went through the images and removed 20 images with major quality issues. For the deep learning approach to tongue and papillae segmentation, a convolutional neural network similar to the U-Net proposed by Ronneberger and colleagues [38] was used. The model’s architecture is shown in **Fig. 6**.

**Fig. 6.**
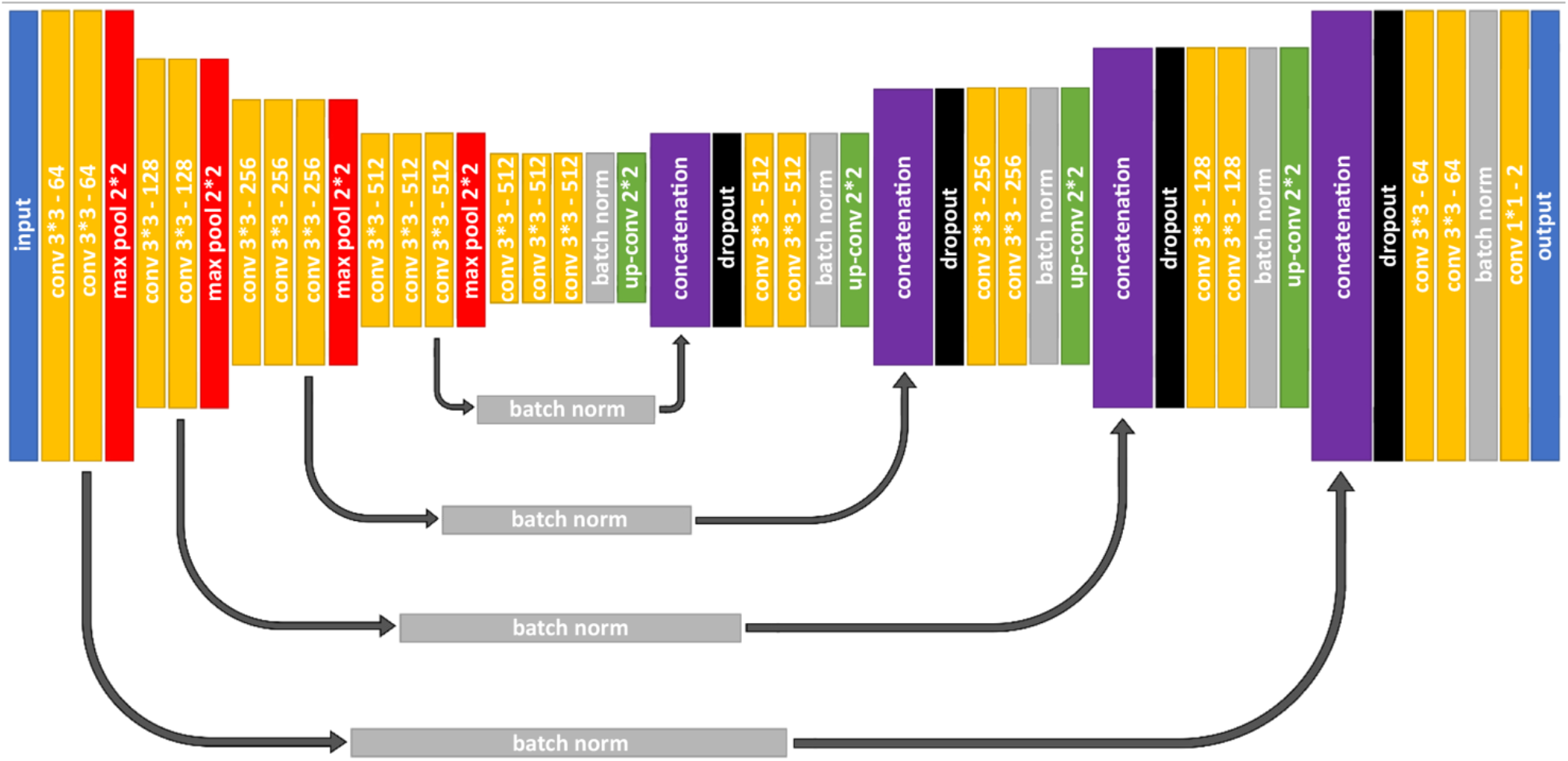
The U-Net based architecture used to segment the tongue and the papillae.

The encoding part of the model is structured identically to a VGG16 [39], but without the dense layers. This allows to load ImageNet weights and apply transfer learning on feature extraction. The rest of the network is modelled after the U-Net, but with added batch normalization and drop-out layers for regularization. An example of the input and target data used in the modelling is shown in **Fig. 7 a-c**.

**Figure 7 a-c:**
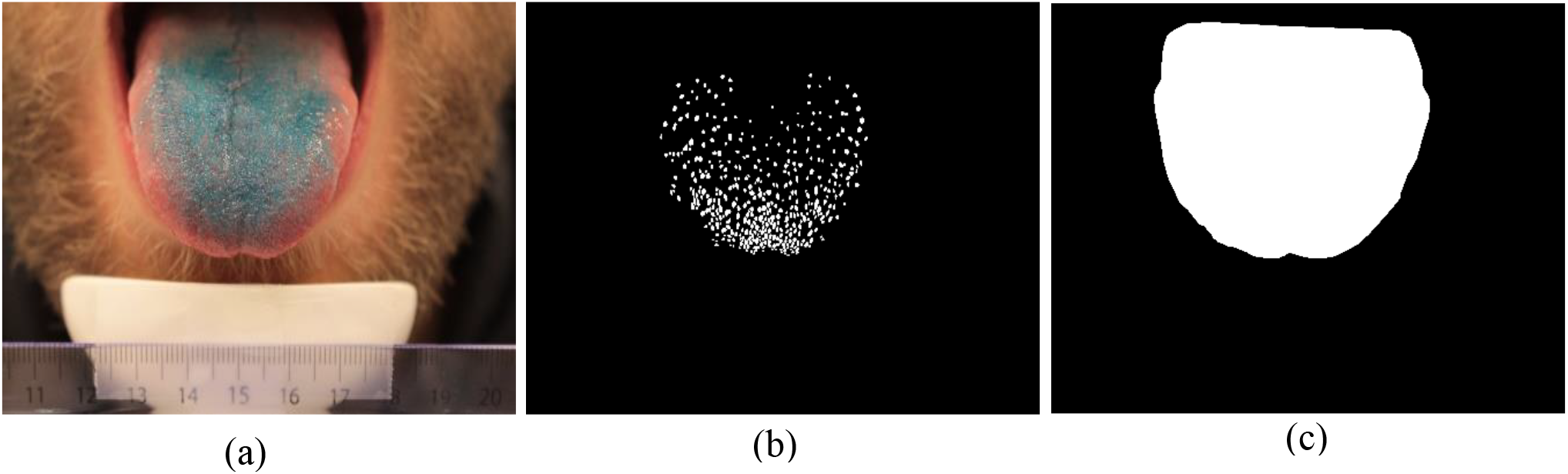
(a) A tongue image, (b) its manual papillae annotation, and (c) its manual tongue annotation.

Considering how densely packed the papillae are and that they only make up a small part of the total image area (TOT), a weighted IoU loss-function was used for network optimization:

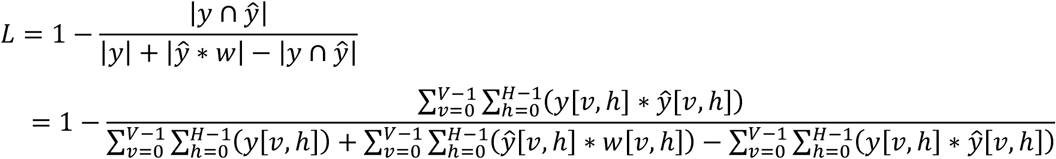

Where ***y, ŷ*** and ***w*** represent the true mask, the predicted mask, and the loss weights respectively. The loss-function penalizes errors in-between neighbouring papillae much harder than errors in other areas to improve object separation. Since the IoU loss is measured in terms of overlap instead of absolute difference, the model is less sensitive to the extreme class imbalance.

For papillae segmentation, the loss weights were computed from the true masks by identifying the pixels that are within a 15 pixels distance from at least two papillae and increasing their weights tenfold (**Fig. 8 a-b**).

**Figure 8 a-b.**
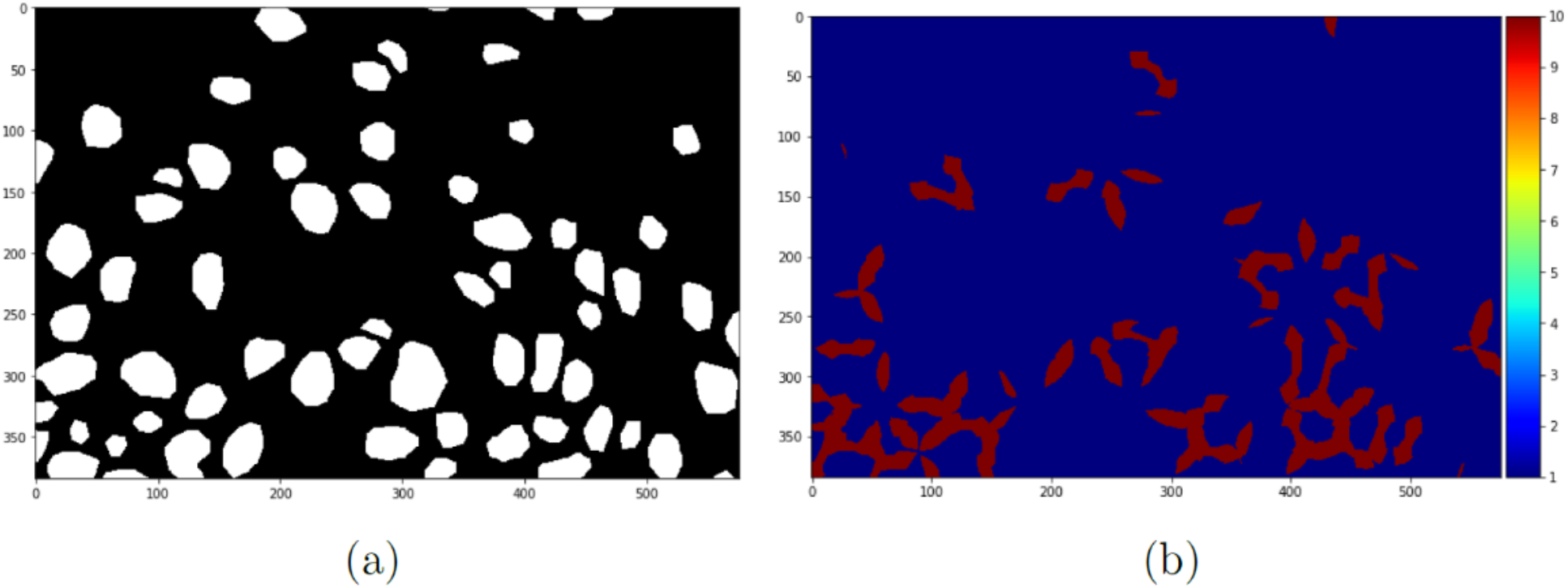
(a) A patch of papillae segmentation and (b) its loss weights.

For tongue segmentation, the same loss-function was used. In this case, the loss weights were drawn by hand and had the purpose of forcing the model to better learn the difference between the tongue and the lip. An example of the loss weights for a tongue is shown in **Fig. 9 (a-b)**. Note that although there are 4 colours, only the lip (green) has its weight increased 5-fold.

**Fig 9 a-b.**
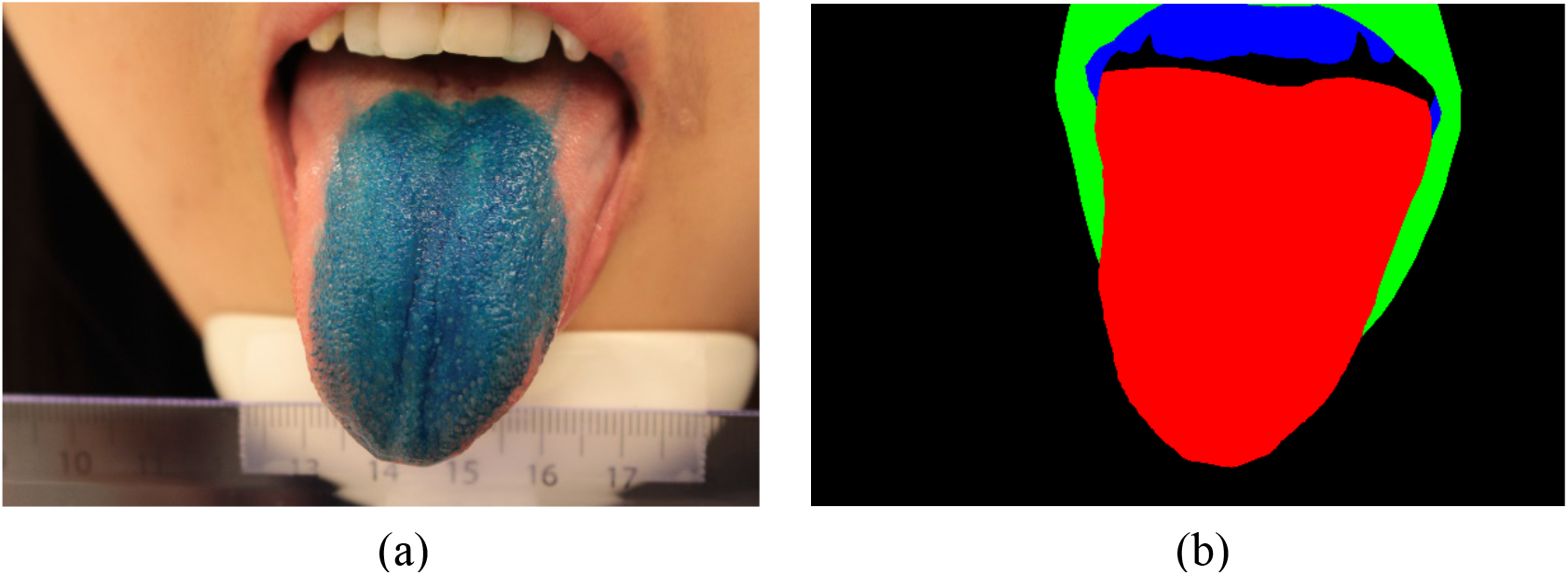
(a) A tongue and (b) its corresponding weights.

In total, 29 selected tongue pictures were used for the training and validation of the papillae segmentation model in a random 8:2 split. Extensive image augmentation was applied to compensate for the relatively small dataset size. The original images along with their masks were cut into patches of 384 x 576 pixels to keep the GPU memory manageable while avoiding information loss. To reduce unnecessary training time, image patches where papillae could not exist, such as regions outside of the tongue, were left out. A sliding window ensemble was used to return output segmentations to their original size at the end.

The training process for tongue segmentation was very similar to that of papillae segmentation, except a slightly bigger training and validation set consisting of 33 tongue pictures was used. Since the tongue took up a much larger proportion of the total area, images and masks were downscaled instead of cutting them into patches.

To achieve accurate counting and measurements, the raw papillae segmentations were first cleaned up using a series of morphological operations. A multiplication was then applied with the corresponding tongue segmentations to remove the false positives outside the tongue. To get rid of the filiform papillae that were erroneously detected as FP, the objects remaining were further filtered by calculating the mean area of the largest 25% of the objects and removing everything smaller than one-fifth of this area. The reason for filtering filiform papillae in this way, instead of thresholding with a specific minimum area, was to account for the fact that the “normal” range of FP area for different individuals were different. The actual papillae counts were extracted by adding up the number of individual objects found with connected components.

### Data Analysis

The normality assumption of the FP density distributions from manual and automated counts was tested by the Shapiro-Wilk W test (α =0.05), and if found to be non-normally distributed, a non-parametric correlation analysis was used. Spearman’s correlation coefficients (ρ_s_) were used to assess the agreement among the different manual and automated counts, with significance defined as p <0.05. The correlation between PROP intensity ratings and FP counts (both manual and automated) was assessed by Spearman’s correlation analysis. Statistical analysis was performed using IBM SPSS statistical software version 25 (SPSS Inc, USA).

The uncertainty of mean FP count was estimated with bootstrapping, where the subset the full taster classes was compared with random sampling with a 1,000 times replacement. The mean FP count was then computed for all 1,000 subsets, resulting in a distribution that could be used to infer where the real value most likely would be.

The statistical significance of a difference in mean FP count between two taster classes was examined using a permutation test, and the power of the permutation test was estimated by bootstrapping: for any pair of taster classes being compared, we performed 1,000 rounds of bootstrapping. At the start of each round, the bootstrapping size was randomly set to 40% ~ 100% of the original class size, and the same percentage was used for both classes to maintain the relative size ratio. A 10,000 round permutation test was then performed, where the difference in mean FP count of the two bootstrapped taster classes against that of two randomly sampled groups were compared. The percentage of times where the random difference was larger than the class difference, also known as the p-value, was recorded before moving to the next bootstrapping round.

In a typical permutation test, a threshold value of 0.05 is used as the significance criterion. This means that if less than 5% of the random differences are larger than the difference by the group, one can reject the null hypothesis that there is no difference between the two groups. With this power estimate, one also has an estimate of the uncertainty of the p-value. Thus, instead, we calculate the frequency sum of p-values larger than 0.10 was calculated. If this sum makes up more than 40% of the total cases, then we would reject the null hypothesis.

## Author Contributions

CC, JL, JS, EP, WLPB designed the study. JL acquired the data. CC, CW, JS performed statistical analysis and data interpretation. CC, CW, JS, WLPB wrote the manuscript. CC, CW, EP, JS, WLPB regularly discussed the experiments, analysed the results, and provided useful suggestion during the project. All authors read and approved the final manuscript.

## References

1. Chandrashekar, J., Hoon, M. A., Ryba, N. J., & Zuker, C. S. The receptors and cells for mammalian taste. Nature. 444(7117), 288 (2006).

2. Miller, I., & Reedy, F. Quantification of fungiform papillae and taste pores in living human subjects. Chem Senses. 15, 281–294 (1990).

3. Miller, I., & Reedy, F. Variations in human taste bud density and taste intensity perception. Physiol Behav. 47, 1213–1219 (1990).

4. Duffy, V. B., et al. Bitter receptor gene (TAS2R38), 6-n-propylthiouracil (PROP) bitterness and alcohol intake. Alcohol.: Clin. Exp. Res. 28(11), 1629–1637 (2004).

5. Essick, G. K., Chopra, A., Guest, S., & McGlone, F. Lingual tactile acuity, taste perception, and the density and diameter of fungiform papillae in female subjects. Physiol Behav. 80, 289–302 (2003).

6. Hayes, J. E. Allelic variation in TAS2R bitter receptor genes associates with variation in sensations from and ingestive behaviors toward common bitter beverages in adults. Chem Senses. 36, 311–319 (2011).

7. Nachtsheim, R., & Schlich, E. The influence of 6-n-propylthiouracil bitterness, fungiform papilla count and saliva flow on the perception of pressure and fat. Food Qual Prefer. 29, 137–145 (2013).

8. Nachtsheim, R., & Schlich, E. The influence of oral phenotypic markers and fat perception on fat intake during a breakfast buffet and in a 4-day food record. Food Qual Prefer. 32, 173–183 (2014).

9. Fischer, M. E., et al. Factors related to fungiform papillae density: The beaver dam offspring study. Chem. Senses. 38(8), 669–677 (2013).

10. Dinnella, C., et al. Individual variation in PROP status, fungiform papillae density, and responsiveness to taste stimuli in a large population sample. Chem. Senses. 43(9), 697–710 (2018).

11. Nuessle, T. M., Garneau, N. L., Sloan, M. M., Santorico, S. A. Denver papillae protocol for objective analysis of fungiform papillae. J. Vis. Exp. 100:e52860 (2015).

12. Sanyal, S., O’Brien, S. M., Hayes, J. E., Feeney, E. L. TongueSim: development of an automated method for rapid assessment of fungiform papillae density for taste research. Chem. Senses. 41, 357–365 (2016).

13. Piochi, M., et al. Comparing manual counting to automated image analysis for the assessment of fungiform papillae density on human tongue. Chem. Senses. 42(7), 553–561 (2017).

14. Segovia, C., Hutchinson, I., Laing, D. G., Jinks, A. L. A quantitative study of fungiform papillae and taste pore density in adults and children. Dev Brain Res. 138:135–146 (2002).

15. Feeney, E. L., & Hayes, J. E. Regional differences in suprathreshold intensity for bitter and umami stimuli. Chemosens Percept. 7, 147–157 (2014).

16. Shahbake, M., Hutchinson, I., Laing, D. G., Jinks, A. L. Rapid quantitative assessment of fungiform papillae density in the human tongue. Brain Res. 1052, 196–201 (2005).

17. Piochi, M., Dinnella, C., Prescott, J., & Monteleone, E.. Associations between human fungiform papillae and responsiveness to oral stimuli: Effects of individual variability, population characteristics, and methods for papillae quantification. Chem. Senses. 43(5), 313–327 (2018).

18. Garneau, N. L., Nuessle, T. M., Sloan, M. M., Santorico, S. A., Coughlin, B. C., & Hayes, J. E. Crowdsourcing taste research: genetic and phenotypic predictors of bitter taste perception as a model. Front Integr Neurosci. 8, 33 (2014).

19. Eldeghaidy, S., et al. An automated method to detect and quantify fungiform papillae in the human tongue: validation and relationship to phenotypical differences in taste perception. Physiol Behav. 184, 226–234 (2017).

20. Valencia, E., et al. Automatic counting of fungiform papillae by shape using cross-correlation. Comput Biol Med. 76, 168–172 (2016).

21. Kraggerud, H., Wold, J. P., Høy, M., & Abrahamsen, R. K. X-ray images for the control of eye formation in cheese. Int J Dairy Technol. 62, 147–153 (2009).

22. Ríos, H. V., Valencia, E., Montes, F. M., Marín, A., Silva, E., & Herrera, R. Recognition of fungiform papillae in tongue images. In CONIELECOMP 2012, 22nd International Conference on Electrical Communications and Computers. 245–247 (2012).

23. Martin, L. J., & Sollars, S. I. Contributory role of sex differences in the variations of gustatory function. J. Neurosci. Res. 95(1-2), 594–603 (2017).

24. Yackinous, C. A., & Guinard, J. X. Relation between PROP (6-n-propylthiou-racil) taster status, taste anatomy and dietary intake measures for young men and women. Appetite. 38, 201–209 (2002).

25. Just, T., Pau, H. W., Witt, M., & Hummel, T. Contact endoscopic comparison of morphology of human fungiform papillae of healthy subjects and patients with transected chorda tympani nerve. Laryngoscope. 116(7), 1216–1222 (2006).

26. Bajec, M. R., & Pickering, G. J. Thermal taste, PROP responsiveness, and perception of oral sensations. Physiol Behav. 95, 581–590 (2008).

27. Bakke, A., & Vickers, Z. Relationships between fungiform papillae density, PROP sensitivity and bread roughness perception. J Texture Stud. 39, 569–581 (2008).

28. Bakke, A., Vickers, Z., Marquart, L., & Sjoberg, S. Consumer acceptance of refined and whole wheat breads. J Food Sci. 72, S473–S480 (2007).

29. Correa, M., Hutchinson, I., Laing, D. G., & Jinks, A. L. Changes in fungiform papillae density during development in humans. Chem Senses. 38, 519–527 (2013).

30. Masi, C., Dinnella, C., Monteleone, E., & Prescott, J. The impact of individual variations in taste sensitivity on coffee perceptions and preferences. Physiol Behav. 138, 219–226 (2015).

31. Piochi, M., et al. Individual variation in fungiform papillae density with different sizes and relevant associations with responsiveness to oral stimuli. Food Qual Prefer. 78, 103729 (2019).

32. Cattaneo, C., Liu, J., Bech, A. C., Pagliarini, E., & Bredie, W. L. Cross-cultural differences in lingual tactile acuity, taste sensitivity phenotypical markers, and preferred oral processing behaviors. Food Qual Prefer. 80, 103803, (2020).

33. Hayes, J. E., Sullivan, B. S., & Duffy, V. B. Explaining variability in sodium intake through oral sensory phenotype, salt sensation and liking. Physiol Behav. 100, 369–380 (2010).

34. Cattaneo, C., et al. New insights into the relationship between taste perception and oral microbiota composition. Sci Rep. 9(1), 1–8, (2019).

35. Prescott, J., Soo, J., Campbell, H., & Roberts, C. Responses of PROP taster groups to variations in sensory qualities within foods and beverages. Physiol Behav. 82, 459–469, (2004).

36. Bartoshuk, L. M., Duffy, V. B., Green, B. G., Hoffman, H. J., Ko, C. W. Valid across-group comparisons with labeled scales: the gLMS versus magnitude matching. Physiol Behav. 82(1), 109–114, (2004).

37. Laureati, M., et al. Associations between food neophobia and responsiveness to “warning” chemosensory sensations in food products in a large population sample. Food Qual Prefer. 68, 113–124 (2018).

38. Ronneberger, O., Fischer, P., & Brox, T. U-net: Convolutional networks for biomedical image segmentation. In International Conference on Medical image computing and computer-assisted intervention. 234–241 (Springer, Cham. 2015)

39. Simonyan, K., & Zisserman, A. Very deep convolutional networks for large-scale image recognition. arXiv preprint arXiv: 1409.1556 (2014).

